# Decision making and best practices for taxonomy-free eDNA metabarcoding in biomonitoring using Hill numbers

**DOI:** 10.1101/2020.03.31.017723

**Authors:** Elvira Mächler, Jean-Claude Walser, Florian Altermatt

## Abstract

Environmental DNA (eDNA) metabarcoding raises expectations for biomonitoring to cover organisms that have hitherto been neglected or excluded. To bypass current limitations in taxonomic assignments due to incomplete or erroneous reference data bases, taxonomic-free approaches are proposed for biomonitoring at the level of operational taxonomic unites (OTUs). However, this is challenging, because OTUs cannot be annotated and directly compared to classically derived data. The application of good stringency treatments to infer validity of OTUs and the clear understanding of the consequences to such treatments is thus especially relevant for biodiversity assessments. We investigated how common practices of stringency filtering affect diversity estimates based on Hill numbers derived from eDNA samples. We collected eDNA at 61 sites across a 740 km^2^ river catchment, reflecting a spatially realistic scenario in biomonitoring. After bioinformatic processing of the data, we studied how different stringency treatments affect conclusions with respect to biodiversity at the catchment and site levels. The applied stringency treatments were based on the consistent appearance of OTUs across filter replicates, a relative abundance cut-off and rarefaction. We detected large differences in diversity estimates when accounting for presence/absence only, such that the detected diversity at the catchment scale differed by an order of magnitude between the treatments. These differences disappeared between the stringency treatments with increasing weighting of the OTUs’ abundances. Our study demonstrated the usefulness of Hill numbers for comparisons between data sets with large differences in diversity, and suggests best practice for data stringency filtering for biomonitoring.

## 1 Introduction

Increasing anthropogenic stressors, such as habitat degradation or pollution, threaten biodiversity and related ecosystem services in freshwater habitats (Dudgeon et al., 2006; Grooten, Almond, et al., 2018; Vörösmarty et al., 2010). To protect and preserve these precious habitats, effective biomonitoring is essential to establish and implement good management practices (e.g., Hering et al., 2006; Johnson, Furse, Hering, & Sandin, 2007; Kelly et al., 2008). Historically, biomonitoring has been restricted to surveys of a few indicator taxa, such as fish, certain groups of benthic macro-invertebrates, and diatoms (Barbour, Gerritsen, Snyder, & Stribling, 1999; Tachet, Richoux, Bournaud, & Usseglio-Polatera, 2010), for which changes in richness and abundance were used to assess the state of ecosystems. Extensive procedures exist for such assessments, often including field collection of specimens and their subsequent identification in the lab. However, the acquisition of adequate biomonitoring data on most taxa is challenging because of time, labor, and cost (Ferraro, Cole, DeBen, & Swartz, 1989; Haase et al., 2004). Environmental DNA (eDNA) analysis and metabarcoding have been proposed as alternative methods with the potential to improve the field of biomonitoring (Bohmann et al., 2014; Deiner et al., 2017; Leese et al., 2018; Pawlowski et al., 2018), because they allow to detect and describe communities at unprecedented scales due to the massive parallel sequencing (Altermatt et al., 2020; Bálint et al., 2018; Taberlet, Coissac, Hajibabaei, & Riesenberg, 2012).

While initial metabarcoding approaches have mostly focused on mirroring or matching data on the well-established indicator taxa (e.g., Douglas et al., 2012; Elbrecht, Vamos, Meissner, Aroviita, & Leese, 2017; Visco et al., 2015), (eDNA) metabarcoding allows us to go well-beyond these indicator groups and has raised the expectation to improve the detection of taxonomic groups that have hitherto been ignored, for example because morphological identification is difficult. Consequentially, eDNA metabarcoding may be offering new opportunities to include currently underused groups of organisms for biomonitoring (e.g., chironomids, Czechowski, Stevens, Madden, & Weinstein, 2020; or oligochaetes, Vivien, Apothéloz-Perret-Gentil, Pawlowski, Werner, & Ferrari, 2019) or even extend to groups that are abundant but currently excluded in aquatic biomonitoring (e.g., rotifers, ciliates). In this context, an accurate taxonomic assignment of eDNA sequences is seen as a crucial but difficult step, because many of these indicator organisms are not completely or inaccurately covered in reference databases (McGee, Robinson, & Hajibabaei, 2019; Weigand et al., 2019). Also, taxonomic names change over time and entries in databases can differ from currently accepted names. As an alternative, the use of taxonomic-free analysis of eDNA has been proposed. This approach is based on the level of operational taxonomic units (OTU), and does not depend/require any taxonomic association (Apothéloz-Perret-Gentil et al., 2017). An OTU is a cluster of sequences grouped by sequence similarity (Blaxter et al., 2005) and is reflecting a pragmatic proxy for species, but is not necessarily in line with classical taxonomy. Taxonomic-free approaches may facilitate rapid implementation of eDNA in biomonitoring. They are, however, challenging because an OTU has no independent verification of any taxonomic unit, unlike taxonomic names assigned to an OTU that can be compared to classically derived data. Using OTUs thus requires good understanding of the data analysis and the steps involved for validation. Of particular importance, especially for biomonitoring and the assessment of biodiversity, is the decision and implementation of stringency treatments to infer validity of OTUs. Multiple strategies to omit OTUs have been reported (see Deiner et al., 2017). Some of these steps are also frequently implemented for eDNA metabarcoding analysis resulting in a taxa list, but the final interpretation of the data is often made in comparison to previously detected taxa in the study area (e.g., Deiner, Fronhofer, Mächler, Walser, & Altermatt, 2016; Hänfling et al., 2016; Mächler et al., 2019). Unfortunately, implemented stringency treatments are often based on somewhat arbitrary decisions but their effects on the downstream diversity analysis has been rarely investigated. Especially in case of taxonomy-free eDNA approaches, we need better understanding of how our decisions during the analysis are affecting the outcome.

After passing stringency selection, OTUs can be used to calculate and compare diversity and richness measures. Virtually all biomonitorings and assessments of biodiversity are based on such data giving numbers or divergence in diversity, and describing diversity at the regional, local or between-site scale (gamma, alpha or beta diversity) to inform decision making processes (e.g., biodiversity monitoring program in Switzerland, BDM Coordination Office, 2009, 2014; or the Multiple Species Inventory and Monitoring for the National forest system in USA, Manley & Van Horne, 2006). While this is common practice, diversity is often calculated in non-compatible ways across its scales or studies, which is severely limiting comparability between the diversity levels, as well as between studies, and thus challenges interpretation and cross-comparisons. Furthermore, it is often neglected that alpha, gamma, and beta diversities are often related in an additive or multiplicative way. Consequently, calculation of beta diversity is not independent from alpha diversity and can lead to incorrect conclusions when comparing beta values of regions with different alpha diversities (Jost, 2007). The framework of Hill numbers (Hill, 1973) has been developed to avoid this problem. Hill numbers mathematically unify the broad array of diversity concepts through incorporating relative abundance and species richness. As a result, all Hill numbers are in the intuitive units of ‘species’, overcome many of the shortcoming of traditionally used diversity metrics and different diversity levels calculated as Hill numbers can be directly compared to each other. While the framework of Hill numbers is already relatively well-established in community ecology (Chao, Chiu, & Jost, 2014; Chase et al., 2018; Jost, 2007), its advantages have only more recently been suggested for DNA-based analysis of micro-organisms (Kang, Rodrigues, Ng, & Gentry, 2016) and macro-organisms (Alberdi & Gilbert, 2019a).

In our study, we investigated how common practices of OTU stringency filtering are affecting diversity estimates based on Hill numbers derived from eDNA samples. We collected eDNA at many sites across a river catchment, reflecting a spatially realistic scenario in riverine biomonitoring. At each site, we collected three replicates of eDNA samples and used a metabarcoding approach by amplifying the cytochrome c oxidase subunit I (COI) region to uncover a broad spectrum of eukaryotic diversity, containing both multiple currently used as well as potential future indicator groups. After bioinformatic data preparation with established quality filtering and clustering criteria, we then studied how different stringency treatments applied to the analysis of the eDNA data affect the interpretation and conclusions with respect to biodiversity: The first set of strategies used were based on consistent presence of each operational taxonomic unit across filter replicates (Alberdi, Aizpurua, Gilbert, & Bohmann, 2018; Beentjes, Speksnijder, Schilthuizen, Schaub, & van der Hoorn, 2018; Mächler et al., 2019). The other strategies were based on a relative abundance cut-off value (Elbrecht & Leese, 2015; Macher et al., 2018; Taberlet, Bonin, Zinger, & Coissac, 2018; Yamamoto et al., 2017) by comparing the data from two different MiSeq runs, and the rarefaction of the data set to the smallest sequencing depth of a sampling site (Taberlet et al., 2018). We investigated how these different selections of stringency treatments are affecting diversity detected at the catchment and the site levels. We conducted all of our analyses in the Hill diversity framework, which not only allowed comparison of these strategies, but also gives direct and intuitive comparability of diversities at different levels, crucial to unify and generalize biomonitoring studies.

## 2 Material and Methods

We used a data set specifically collected for the analysis of diversity patterns across a large riverine ecosystem, as commonly surveyed in biomonitoring studies. We investigated how diversity estimates derived by eDNA are affected by stringency decisions during analytical steps after the bioinformatic data preparation. Below, we first describe the collection of the data set and second how we applied stringency decisions during the data analysis.

### 2.1 Data set collection

We used data on diversity of eukaryotes estimated by eDNA metabarcoding in a riverine network. To do so, we sampled eDNA at 61 sites in a 740 km^2^ catchment of the river Thur (Fig. 1A), northeastern Switzerland, in June 2016. The streams at the selected sites range from first to seventh stream order and cover an elevation gradient from 472 to 1241 m above sea level. All details of collection and preparation of the herein presented data are extensively described in a previous study (Mächler et al., 2019), which analysed the diversity of the taxonomic subgroups of may-, stone-, and caddisflies. Here we give only a short overview of the general field and laboratory methods, necessary for the understanding of the data and the experimental setup (Fig. 1B), and we then explain the analysis of the complete metabarcoding data set covering all eukaryotic eDNA.

**Figure 1.**
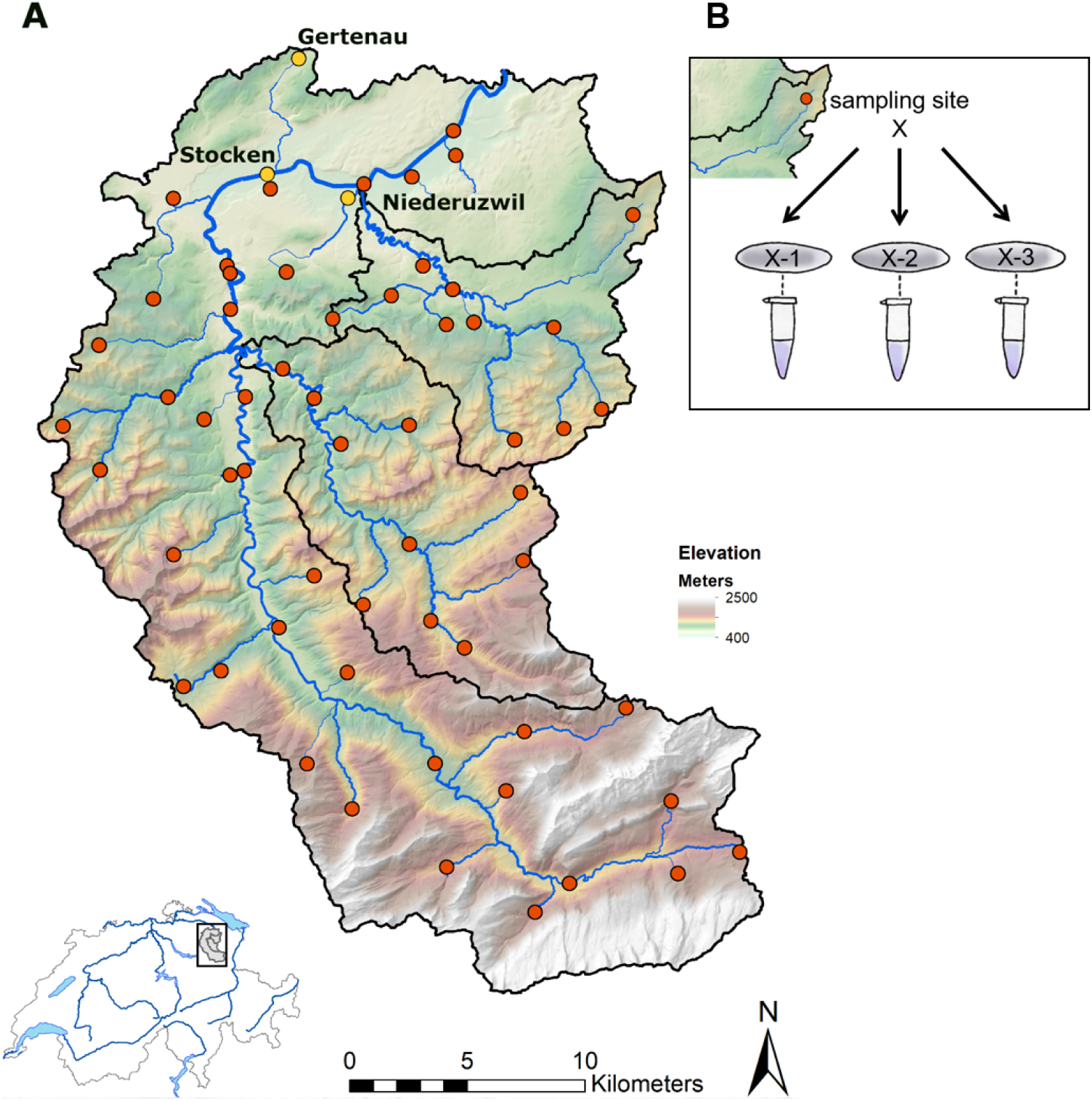
Overview of the study catchment (A) and the 61 individual sampling sites indicated with red dots. Yellow points show the three sites that we picked specifically for further site analysis (see Fig. 5). At each site (B), we collected three individual and independent filter replicates that were separately extracted and tagged. Data source: swisstopo: VECTOR200 (2017), DEM25 (2003), SWISSTLM3D (2018); BAFU: EZG (2012); Bundesamt für Landestopographie (Art.30 Geo IV): 5,704,000,000, reproduced by permission of swisstopo/JA100119.

At each site, we filtered 250 mL of water on a single GF/F glass fiber filter (25 mm diameter, 0.7 µm pore size, Whatman International Ltd., Maidstone, U.K.). We replicated the filtration three times (Fig. 1B). After filtration, we transferred the filters to independent Eppendorf tubes and stored them immediately in the dark on ice. At the end of the sampling day, samples were moved to –20 °C until extraction. In total, we collected 183 filter samples over the whole sampling campaign. At the beginning of each field day, we produced replicated filter controls (FC) consisting of 250 mL of UVC light treated nanopore water, to estimate possible contamination of used material. Over the 11 sampling days, we produced 33 filter controls. Additional to the eDNA samples, we measured environmental variables (pH, conductivity, organic phosphor) and calculated mean temperature based on hourly measurements of Hobo pendant logger (model UA-001-08; Onset Computer Corporation, Pocaset, MA) over the course of five weeks at each site.

The extraction was performed in a specialised clean-lab facility (Eawag, Switzerland; see Deiner, Walser, Mächler, & Altermatt, 2015; Mächler, Deiner, Steinmann, & Altermatt, 2014) with the DNeasy Blood & Tissue kit (Qiagen GmbH, Hilden, Germany) following an adjusted spin-column protocol. We eluted the filter samples in 75 µL of AE buffer. On each batch of extraction, we included an extraction control (EC), resulting in eight ECs. To remove inhibition of the field samples, we cleaned all extractions with OneStep PCR inhibitor removal kit (Zymo Research, Irvine, California). We used the Illumina MiSeq dual-barcoded 2-step PCR amplicon sequencing approach to sequence a 313 base pair fragment of the COI barcoding region (Geller, Meyer, Parker, & Hawk, 2013; Leray et al., 2013). We performed the first PCR with primers containing an Illumina adaptor-specific tail, a heterogeneity spacer and the amplicon target site. This PCR reaction consisted of 25 µL reaction mix and 5 µL of eDNA template, with the exception of samples coming from five sites (Site ID 8, 18, 16, 32 and 47) where we used 5 µL of a 1:10 dilution of the eDNA sample, due to amplification difficulties of pure eDNA. A negative control (NC), consisting of 5 µL sigma water, and a positive control (PC), consisting of 1 µL artificially produced dummy DNA (Mächler et al., 2019) and 4 µL of a randomly selected eDNA sample, were run along each 96-well PCR plate, resulting in a total of 3 NCs and 3 PCs. We produced three PCR replicates of each sample and pooled them before cleaning up with SPRI beads (Applied Biological Materials Inc., Richmond, Canada). The second PCR consisted of 25 µL reaction mix, 5 µL Index N, 5 µL Index S (Nextera XT Index kit v2) and 15 µL of cleaned template. We used 10 PCR cycles to index the samples; thereby the filter replicates were tagged individually to allow analysis between the filter replicates. We then cleaned 25 µL of the indexed reaction with SPRI beads and quantified the clean samples with the Spark 10M Multimode Microplate Reader (Tecan^®^ Group Ltd., Männedorf, Switzerland). We pooled the samples equimolar and cleaned the resulting pool again with SPRI beads. We added 16 pM of libraries and 16 pM of 10% PhiX control to the flow cell. On an Illumina MiSeq platform, we performed a paired-end 600 cycle (2 × 300 nt) sequencing with the Reagent Kit v3 (300 cycles). We conducted a second run on the identical Illumina MiSeq machine with the same libraries, where we added again 16 pM of each PhiX control and the pooled libraries.

The NGS data was demultiplexed with the MiSeq Reporter and the read quality was checked with FastQC (Andrews, 2015). Next, raw reads were end-trimmed (usearch, version 10.0.240) and merged (Flash, version 1.2.11). We then removed the primer sites (cutadapt, version 1.12) and quality-filtered the data (prinseq-lite, version 0.20.4). We used UNOISE3 (usearch, version 10.0.240) to determine zero-radius OTUs (ZOTUs). ZOTUs are valid operational taxonomic units that are corrected for point errors to retrieve reliable sequence clustering and are further filtered to remove chimeric sequences (Edgar, 2016). To reduce sequence diversity, we conducted an additional clustering at 99% sequence identity. To ensure an intact open reading frame, we checked the resulting ZOTUs for stop codons using the invertebrate mitochondrial code.

We cleaned the data by following the description of Evans et al. (2017) due to indication of contamination (i.e., presence of ZOTUs in negative controls). We calculated the relative frequency of each ZOTU appearing in the negative controls by dividing the total reads of an individual ZOTU in all negative controls by the total number of reads in the negative controls. We removed all ZOTUs that occurred with a lower relative frequency within a given field sample. For all further analysis, we removed all samples from the sites that used diluted eDNA for the first PCR (Site ID 8, 16, 18, 23 and 47), as we observed low consistency among the three filter replicates. To obtain consistent results between replicates, it is not recommended to dilute eDNA, even though this is a favored method implemented to reduce inhibition (McKee, Spear, & Pierson, 2015). Fifty-six sampling sites remained and were used for the subsequent analyses.

### 2.2 Hill numbers and stringency treatments

We conducted all our diversity analyses in the framework of Hill numbers, which mathematically unifies diversity concepts based on relative abundances (Hill, 1973; Jost, 2007). As such, the sensitivity of Hill numbers towards abundant taxa can be modulated with the parameter *q*, also known as the order of Hill numbers. A Hill number with an order 0 (*q* = 0) is insensitive to abundance, and is analogue to the classical species richness measure. A Hill number of order 1 (*q* = 1) takes the exact relative abundance into account and is analogue to the exponential of Shannon diversity as alpha diversity measure. A Hill number of order 2 (*q* = 2) gives abundant taxa more weight and is analogue to the inverse of the Simpson index. The partitioning of biological diversities into the classical alpha, beta and gamma diversity can be directly computed and expressed as Hill numbers, whereby alpha measures reflect the average diversity of the sampling sites, gamma the overall diversity, and beta the dissimilarity between sites and the overall diversity, for example of a catchment (Jost, 2007). All Hill numbers are calculated and interpreted as units of ‘species’ (i.e., in our case as units of ZOTUs), can be directly compared across orders (Alberdi & Gilbert, 2019a; Chao et al., 2014; Jost, 2007), and directly allow us to study differences between the stringency treatments.

We then applied five different stringency treatments to identify how these affect diversity patterns in the eDNA data, and thus may directly affect interpretation of biomonitoring and biodiversity-assessments. The first three strategies are based on consistent appearance of ZOTUs across replicates (Alberdi et al., 2018; Beentjes et al., 2018; Mächler et al., 2019). We assessed how often each individual ZOTU was detected across the three filter replicates of an individual site and differentiated between *additive* (counting all ZOTUs found in any of the three replicates), *relaxed* (counting only ZOTUs with a minimal presence in two replicates), and *strict* (counting only ZOTUs with presence in all three replicates). These categories are based on the description of Alberdi et al. (2018), but here we are referring to filter replicates rather than PCR replicates (as in Alberdi et al., 2018). The fourth strategy was based on a relative abundance cut-off values (Elbrecht & Leese, 2015; Macher et al., 2018; Taberlet et al., 2018; Yamamoto et al., 2017). We plotted for each ZOTU its relative abundance in the first run against its relative abundance in the second run (Fig. S1) and identified a valid threshold of 0.005%, as abundances between the two runs were relatively similar above this value and it also corresponds to the recommended percentage to use for studies without the positive control of a mock community (Bokulich et al., 2013). We subsequently refer to this stringency strategy as the *threshold* treatment. As a fifth stringency treatment, we rarefied the data set to the smallest sequencing depth of a sampling site (251,207 reads, all three replicates combined), which we subsequently refer to as the *rarefaction* treatment. These five stringency treatments resulted in five distinct data sets, that were analyzed subsequently in R (R Core Team, 2018, version 3.5.2) and the package ‘phyloseq’ (McMurdie & Holmes, 2013, version 1.24.2).

#### 2.2.1 At catchment level

We were interested in how the different stringency measures are affecting the partitioning of diversity measures at the catchment level. We partitioned for alpha, gamma and beta diversity based on qualitative (i.e., presence/absence, order 0) and quantitative (i.e., abundance based, from order 0.5–2, in steps of 0.5) Hill numbers for each of the five stringency treatments (function div_part in the ‘hilldiv’ R package, Alberdi & Gilbert, 2019b, version 1.5.0). As diversity measures tend to increase with increasing sampling effort, ecologists implement rarefaction to correct for this bias (Gotelli, 2001; Gotelli & Colwell, 2011). Classically, rarefaction is based on sample size, which can be represented by the number of collected individuals (e.g., Simberloff, 1978), or the number of sites visited (e.g., Gotelli, 2001). Alternatively, Good (1953, 2000) suggested a coverage-based approach, which standardizes samples based on their recovered completeness (i.e., the recovered proportion of the total number of individuals and species of a community represented in the sample). We calculated site- and coverage-based rarefaction and extrapolation curves (Chao & Jost, 2012; Colwell et al., 2012) for the catchment diversity with the ‘iNEXT’ R package (Hsieh, Ma, & Chao, 2016, version 2.0.20). To do so, we utilized incidence data, which are based on relative detection (presence/absence) over the whole catchment rather than relative abundances at the individual sites (Alberdi & Gilbert, 2019a; Colwell & Coddington, 1994).

#### 2.2.2 At site level

To assess in how the different stringency treatments are affecting the diversity measures at the individual sampling sites, we calculated Hill numbers for each of the five treatments using presence/absence data (order 0) and abundance based data (order 1 and 2). To identify whether there were significant differences in detected diversity between the stringency criteria or not, we performed the following tests: Within each order, we tested whether the variance is equal between the stringency treatments by using a Bartlett test. When significant, we performed a Kruskal-Wallis test to identify whether there was at least one difference in means. If so, we then did a multiple mean comparison post-hoc test based on rank sums (R package ‘pgirmess’, Giraudoux, 2018, version 1.6.9). We further investigated beta diversity between the individual sites for the orders 0, 1, and 2 (function ‘part_div’, Sørensen-type overlap from the R package ‘hilldiv’, Alberdi & Gilbert, 2019b) and tested for differences between the mean of the stringency treatments as explained above.

To identify how decisions and implementations of stringency criteria are affecting the diversity at individual sites, we selected three different sampling sites (Gertenau, Niederuzwil, and Stocken) from the catchment as proof-of-concept examples. These sites correspond to different stream orders (1, 4, and 7 respectively) and reflect distinct habitats, but are geographically still relatively close to each other, and thus do not experience different large-scale environmental drivers. We investigated sample-size based rarefaction/extrapolation curves with abundance data of the individual sampling sites.

## 3 Results

The two Illumina MiSeq runs resulted in 17,495,861 and 21,026,108 raw sequences, respectively. After the bioinformatic data preparation, 12,619,152 and 14,735,110 merged, primer-trimmed, and quality filtered sequences remained for run 1 and run 2, respectively. In the combined data set, in total 26,544,120 counts assigned to the samples remained for our analysis after the correction for ZOTUs in negative controls. After the application of the five stringency treatments, the resulting five data sets differed mainly by the number of ZOTUs. The biggest difference in numbers of ZOTUs occurred between the *threshold* treatment and the *additive* stringency treatment, with the former containing only 12.4% of the ZOTUs found in the latter (Table 1). Differences in counts were relatively moderate. We detected the biggest difference between the *additive* and *rarefied* treatments, where the *rarefied* treatment had about 43% less counts. The rarefaction curves were mostly saturating for sampling sites of the *relaxed, strict*, and *rarefied* treatments (Fig. S2 B, C, E), but not for most of the sites of the *additive* and *threshold* treatment (Fig. S2 A, D).

**Table 1:** Numbers of reads and ZOTUs for the five stringency measures after cleaning the data set from contamination.

|  | **Additive** | **Relaxed** | **Strict** | **Threshold** | **Rarefied** |
| --- | --- | --- | --- | --- | --- |
| Reads | 24,640,037 | 22,105,491 | 19,401,888 | 22,917,814 | 14,067,592 |
| ZOTUs | 11,129 | 5,948 | 3,167 | 1,385 | 11,102 |

### 3.1 At the catchment level

The partitioning of the five stringency-filtered data sets into Hill numbers of order 0 resulted in alpha and gamma diversity (corresponding to richness) varying over nearly an order of magnitude (Fig. 2, Table S1). The beta diversity estimated by Hill numbers quantifies how many times the diversity of the entire catchment is richer than the average sampling site and varies greatly between the stringency treatments. Beta values indicated that the difference between sites and the catchment data set was largest for the *strict* data set, where the regional (gamma) diversity was nearly 12 times higher than the average local (alpha) diversity, indicating large differences between the diversity at the individual sites. Contrary, the richness for *threshold* treatment was only 2.39 times higher for the catchment than the sites, indicating less differences between the richness of the individual sites.

**Figure 2.**
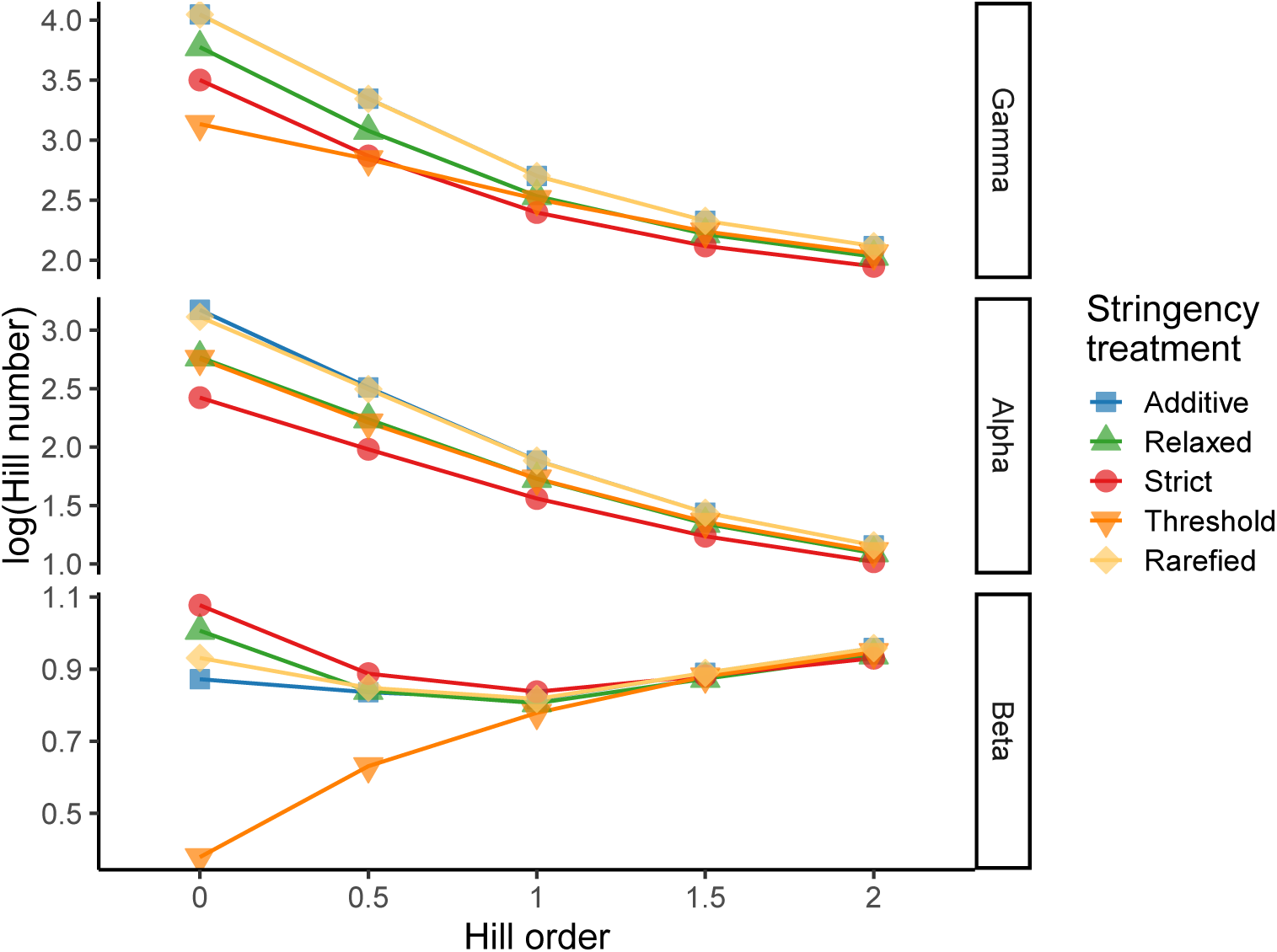
Diversity partitioning of the five stringency treated data sets for Hill numbers of different orders. The higher the order, the more weight is put on abundant ZOTUs; order *q* = 0 reflects presence-absence data. While gamma and alpha diversity behave similar for all the treatments, beta dissimilarity shows differences: For each stringency treatment, the differentiation between the individual sites (alpha) and the overall (gamma) increases with higher weighting of abundances. Hill numbers are presented on a logarithmic scale for better visualization. Exact numbers can be found in Table S1. The colors represent the different stringency treatments: *additive* (blue), *relaxed* (green), *strict* (red), *threshold* (orange), and *rarefied* (yellow).

Taking abundances into account (order 0.5 and higher), the partitioning values for all five data sets became more similar when abundances were stronger weighted, indicating that all stringency data sets were dominated by a few, very abundant ZOTUs (Fig. 2). All five stringency treatments ended up with a beta diversity value of around 9 for order 2 (Table S1). The beta value for the *threshold* data set was increasing from order 0 to order 2, demonstrating that the ZOTUs of this data set were present in many samples and dissimilarity between sites increased with abundance weighting.

Whether the accumulation of diversity with number of sampling sites was saturating or not depends on the used order of Hill numbers (Fig. 3A). For the order *q* = 0, only the *threshold* data set was saturating (even at a very low number of sampling sites), while the other stringency treatments did not reach a saturation phase. With increasing order, the shape of the curves started to saturate with lower number of samples and for the order *q* = 2 all stringency treatments were saturating.

**Figure 3.**
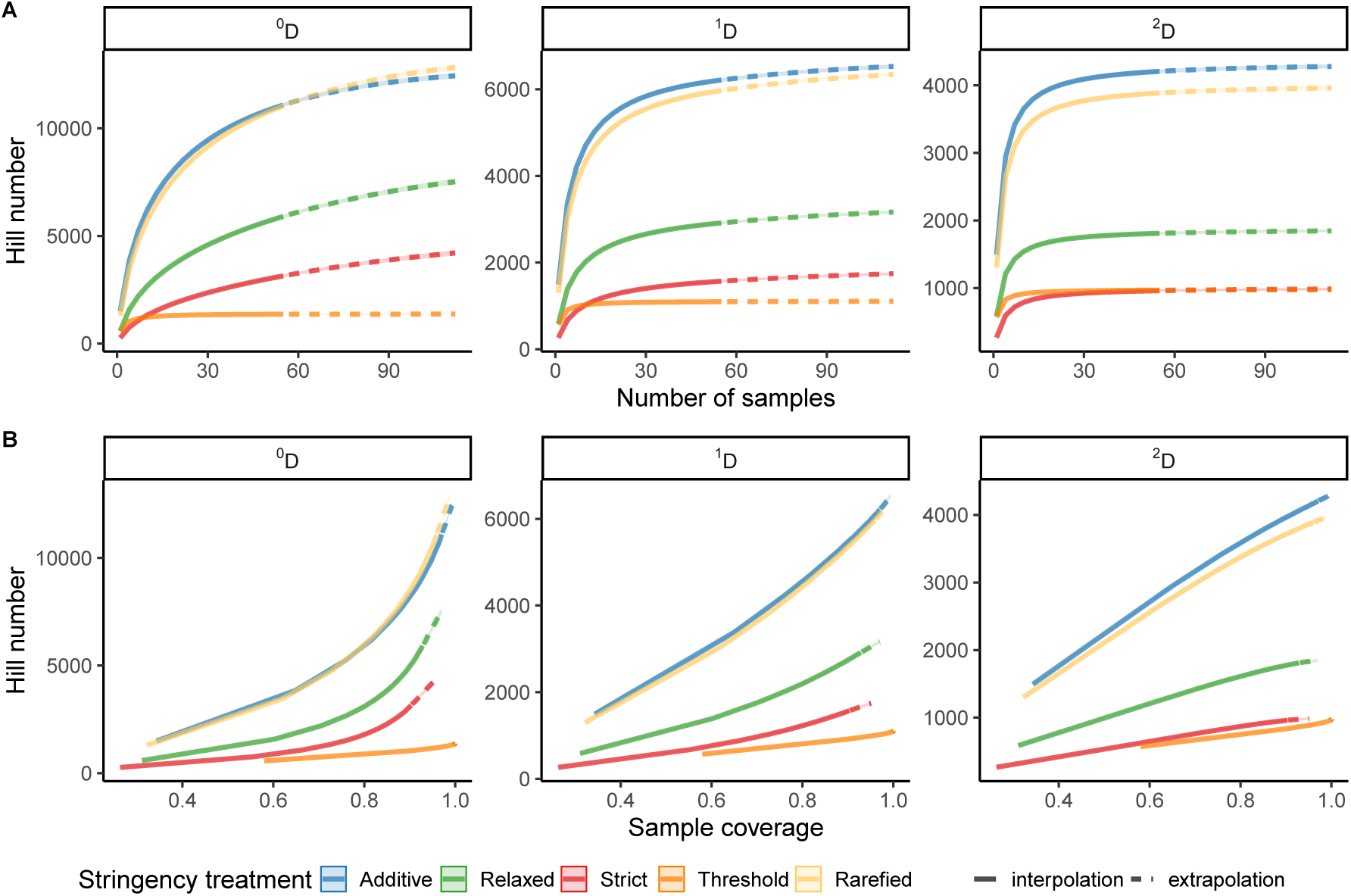
Accumulation curves of Hill diversity based on number of samples (A) and sample coverage (B) over the whole catchment. Diversity for the Hill numbers are calculated for the order *q* = 0 (left panels), *q* = 1 (middle panels) and *q* = 2 (right panels). The solid lines indicate parameter space that is established by rarefaction of the measured data, while dashed lines indicate extrapolated values.

The sample coverage of the catchment diversity was highest for the *threshold* treatment (99.9%), followed by the *additive* (97.1%), *rarefied* (95.6%), *relaxed* (93.0%) and *strict* with a sample coverage of 90.3% (Fig. 3B). The coverage-based rarefaction curves turned towards a positive linear relationship with increasing order of Hill numbers, revealing that with higher abundance weighting the coverage increases only linearly with each additional sample.

### 3.2 At sampling site level

In biomonitoring studies, the diversity uncovered at the site level is important, for example, to define management priorities between individual sites. When analysing the mean of the alpha diversity, no significant difference were ever observed for the *additive* and *rarefied* stringency criteria, no matter how strong abundances were weighted (Table 2, Fig 4). Additionally, these two stringency treatments were always significantly different from the *strict* treatment. When comparing the mean of the recovered diversities, we observed a dependency on the order of the Hill number (Table 2). The significant difference of the *relaxed* and *threshold* treatment to others was depending on the Hill order. With *q* = 0, both treatments were significantly different to the other treatments, but not each other. With *q* = 1, the *threshold* was significantly different from the *strict*, but the *relaxed* stringency treatment was not different from any other treatment. For *q* = 2, both treatments were indifferent from the other treatments. For the beta diversity between sites we also observed different patterns for the various Hill orders. When looking at *q* = 0 (i.e., Jaccard dissimilarity), all five stringency treatments had significantly different means (Table S2, Fig. S3). With *q* = 1, three different groups appeared: *additive, relaxed*, and *rarefied* were different from *strict*, as well as from the *threshold* treatment, but did not differ from each other, and the *strict* treatment significantly differed from the *threshold* treatment. For the order *q* = 2 (i.e., the Morisita-Horn index) there was no significant difference in means observed.

**Table 2:** Results of Kruskal-Wallis test on ranks of the alpha diversity and the following multiple mean comparisons post-hoc tests based on rank sums. Chi-Squared, degrees of freedom (DF), number of data points (n) and p-values are given. Stringency treatments with the same letter are not significantly different according to multiple comparison post-hoc test after Kruskal-Wallis (p-value = 0.05).

| Kruskal-Wallis test on ranks |  |  |  |  | Multiple mean comparisons |  |  |  |  |
| --- | --- | --- | --- | --- | --- | --- | --- | --- | --- |
|  |  |  |  |  | post-hoc tests |  |  |  |  |
| Hill order | Chi-Squared | DF | n | p-value | Additive | Relaxed | Strict | Threshold | Rarefied |
| 0 | 224.54 | 4 | 280 | < 0.001 | a | b | c | b | a |
| 1 | 39.66 | 4 | 280 | < 0.001 | a | ab | b | a | a |
| 2 | 10.26 | 4 | 280 | 0.0113 | a | ab | b | ab | a |

**Figure 4.**
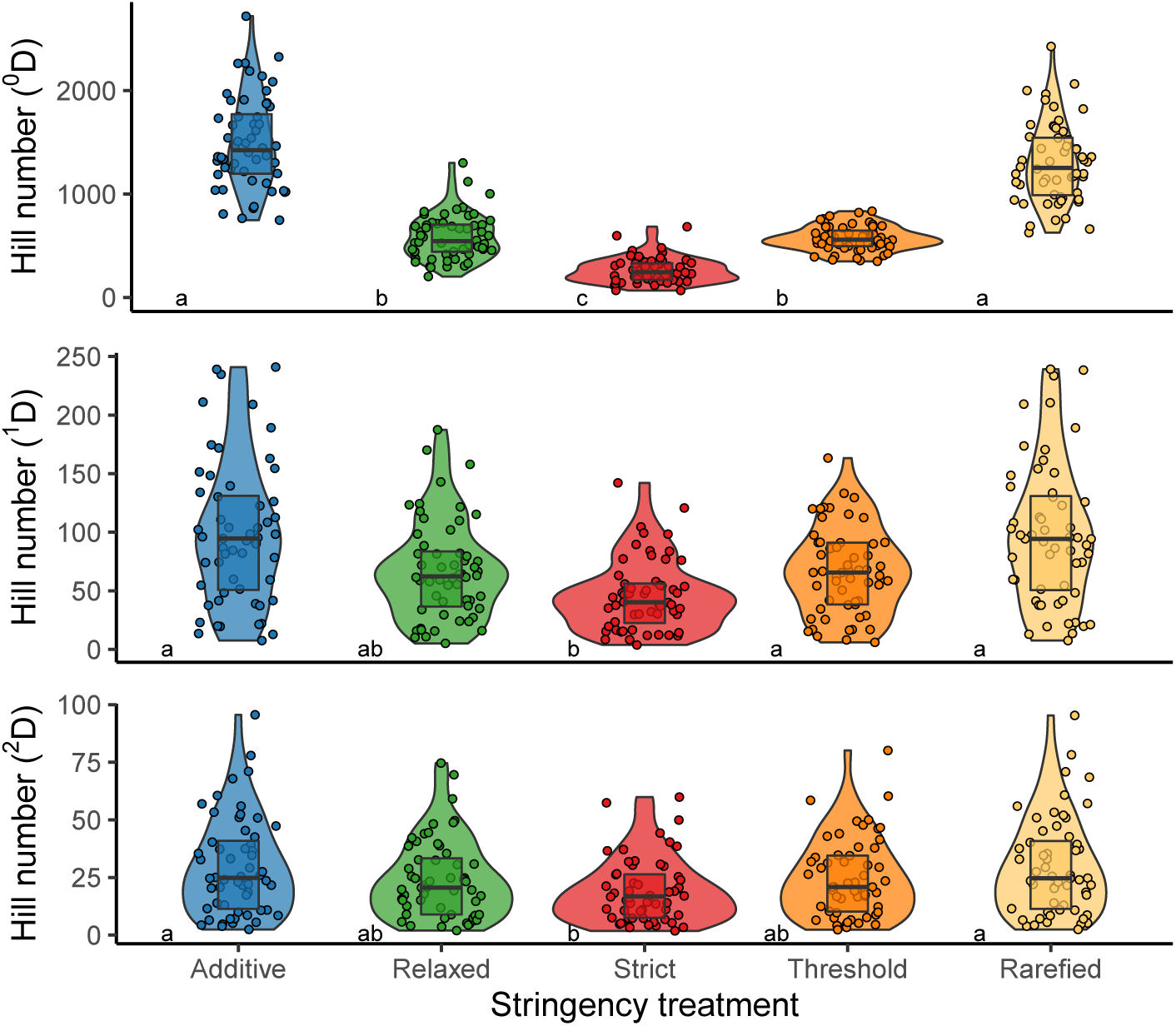
Hill numbers given for the five stringency measures (*additive, relaxed, strict, threshold*, and *rarefied*) and the different orders of Hill numbers (indicated by the exponent of D given in the y-axis). Violin plots depict the distribution of values of the 56 sites. The inserted boxplots are indicating the 25%, 50% (bold) and 75% quantile, respectively. Stringency treatments with dissimilar letters (a, b, and c) are significantly different according to multiple mean comparison test after Kruskal-Wallis applied separately to the individual Hill number orders.

Sequence-based rarefaction curves behaved similarly between the three sites, but differed between orders of Hill numbers (Fig. 5). For the order *q* = 0, the *additive* and *rarefied* stringency treatments did not saturate, while the other three treatments did. However, for orders *q* > 0 the saturation for all five stringency treatments occurred even at a low number of sequences. Surprisingly, the *strict* stringency treatment resulted in highest alpha diversity for higher Hill orders *q* > 0 for the site of Niederuzwil (Fig. 5, middle panel). Increased alpha diversity can be observed if few, but very abundant ZOTUs are leading to low Hill numbers for order *q* > 0. However, without these ZOTUs present in all three samples, the abundance of the remaining ZOTUs are more equally distributed and resulted in a higher diversity even for more stringent criteria.

**Figure 5.**
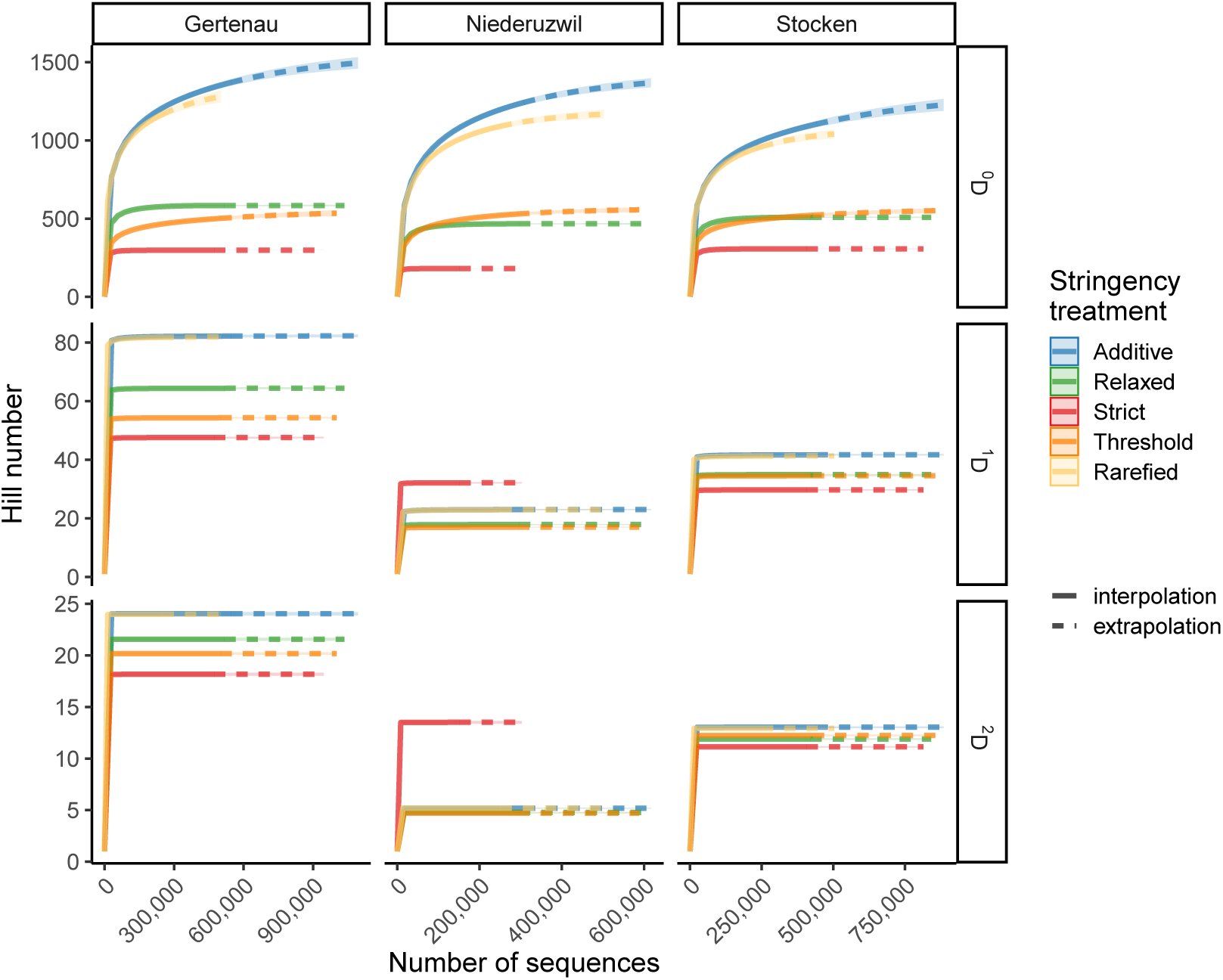
Rarefaction curves of Hill diversity based on number of retrieved sequences for three sampling sites: Gertenau (stream order 1, left panels), Niederuzwil (stream order 4, middle panels), and Stocken (stream order 7, right panels). Diversity for the Hill numbers are calculated for the order *q* = 0 (upper panels), *q* = 1 (middle panels) and *q* = 2 (lower panels). The solid lines indicate parameter space that is established by rarefaction of the measured data, while dashed lines correspond to extrapolated values.

## 4 Discussion

The recent increase in the proposition and use of eDNA metabarcoding for biomonitoring requires that methodological steps in data analysis and calculation of diversity measures are well-understood. This is relevant for metabarcoding data focusing on well-defined taxonomic groups paralleling existing monitorings (e.g., diatoms, Visco et al., 2015; or benthic macroinvertebrates, Elbrecht et al., 2017), but even more so when the metabarcoding extends beyond these groups and is performed in a taxonomy-free approach (Apothéloz-Perret-Gentil et al., 2017; Porter & Hajibabaei, 2018). Biomonitoring goes beyond presence-absence measures, and especially in the context of ecological functions and services provided by an ecosystems, relative or absolute abundances become important. Thus, a coherent understanding of diversity measures needs to consider also abundance-based estimates, and do so in ways that are comparable across and within data sets. This is possible in the mathematical unifying framework of Hill numbers when calculating diversity measures (Alberdi & Gilbert, 2019a; Chao et al., 2014; Jost, 2007).

In this study, we investigated how common practices of stringency filtering in the analysis of eDNA metabarcoding data are affecting diversity estimates. The evaluation of different stringency treatments within the Hill number framework showed that differences between treatments scale with the abundance weighting of the ZOTUs. While we observed large differences between the individual stringency treatments for orders *q* = 0, the differences decreased or even disappeared with the stronger weighting of abundances. Overall, the stringency treatment of *rarefied* and *additive* were leading to very similar results for all diversity measures, potentially due to relatively high sequencing depth for the individual sites minimizing the effect of rarefaction. The *relaxed* treatment produced intermediate estimates of diversity, while the *strict* and *threshold* were usually resulting in the lowest estimates for alpha and gamma diversity. The effects of such stringency treatments on diversity measures have been rarely investigated, but are important to infer validity of (Z)OTUs in the statistical data analysis of taxonomy-free eDNA metabarcoding data.

The applied stringency measures already showed an effect in the overall number of ZOTUs. The consistent presence in at least two (i.e., *relaxed*) or three (i.e., *strict*) replicates reduced the numbers of ZOTUs to half or even a third, respectively. Such a variation in detectability across replicates might be due to stochasticity in the field (i.e., due to heterogeneous distribution of eDNA, Adrian-Kalchhauser & Burkhardt-Holm, 2016; Macher & Leese, 2017), or in the lab through sub-sampling and stochasticity in the PCR reaction. Especially errors or biases in PCR reactions can contribute to an inconsistency across true replicates (Alberdi et al., 2018; Leray & Knowlton, 2017; Murray, Coghlan, & Bunce, 2015), which cannot be eliminated during bioinformatic data preparation. The use of technical (e.g., PCR reactions, Alberdi et al., 2018; Beentjes et al., 2018) or biological (e.g., Macher & Leese, 2017; Mächler et al., 2019) replicates allows to apply such stringency treatments that are capable to infer validity of ZOTUs. Our study highlights, consistent with other studies, the importance of taking multiple replicates, and to use these replicates to get more coherent and robust conclusions on the diversity assessed (Alberdi et al., 2018; Leray & Knowlton, 2017).

The application of a customized abundance thresholds requires either the performance of at least two separate sequencing runs or the implementation of a mock community (Bokulich et al., 2013), with the latter being complicated for complex and broad taxonomic communities. As shown in our study, the two sequencing runs on the same data set allow the identification of good estimates for a cut-off values of minimal read numbers to be included and should be preferred over the selection of a random threshold. We acknowledge that there are higher costs associated with additional sequencing runs, which may be limiting in some monitoring programs. However, in the long run, monitoring programs are only valuable when they provide reliable data, and thus these additional costs may be well-invested. Alternatively, rarefaction can be applied without further investment in replication of samples or runs, but is often heavily criticized (McMurdie & Holmes, 2014), as it tends to loose rare, but potentially essential, ZOTUs that can reflect important aspects of communities (Ainsworth et al., 2015; Eren et al., 2015; Zhan et al., 2013).

Our analysis showed that differences, and thus conclusions on diversity, between the five stringency treatments at the catchment level are large, given that the diversity estimates between treatments spanned nearly an order of magnitude when considering richness only. Also, the differentiation between sites and the overall catchment was limited for the *threshold* treatment for richness, indicating that remaining ZOTUs of the *threshold* stringency treatment had a high prevalence in all the sampling sites, also reflected in the fast saturation of the catchment diversity with accumulating number of samples. However, the individual sites became more distinct with increasing abundance weighting (i.e., most ZOTUs are present in all sampling sites but differ in abundances), an effect that was similar for all stringency treatments for the order *q* ≥ 1. The observed effect might be a specific feature of our study, where we expected little differences between the sites as they are connected and do not have substantial gradients in habitat quality between sites. However, such a data set can reflect a biomonitoring study and it is important to be aware that the application of cut-off values for abundances and the simultaneous use of richness measurements can minimize differences between sites.

When comparing differences between the treatments at the site level, both alpha diversity and beta dissimilarity estimates get more similar with increased weighting of abundances. Nevertheless, the mean alpha diversity of the *strict* treatment was always significantly lower compared to the *additive* and *rarefied* treatments, regardless how strong ZOTU abundances were weighted. But the situation can differ when focusing on an individual site, as shown in one of the selected sampling location of Niederuzwil (Fig. 5).

We generally could observe that differences between the treatments disappeared with higher weighting of abundances. This indicates that weighing abundances may be one way to make data sets more robust when comparing across studies. However, it is yet unclear whether the focus on the abundant (z)OTUs only is an eligible option for biomonitoring studies. Recent research showed that, for classical biomonitoring studies and subsequently derived indices on ecological state, presence-absence data of classical approaches are resulting mostly in the same conclusions then when including abundance information (Beentjes et al., 2018; Buchner et al., 2019). In contrast, our study shows that the eDNA metabarcoding results will differ to quite some extent depending on the used stringency treatment in the data analysis, and thus needs critical considerations of the selected criteria and may be most robust when including information on abundance data. This seemingly contradictory outcome may be explained because indices on the biological state commonly used in ecotoxicology and environmental sciences are already strong integration of complex data, and this integration of multiple information may make them more robust. In our approach the values of interest are much closer to the units ecologists and biodiversity scientists are using, namely number and differentiation of taxa, and those are indeed sensitive to the cut-offs and stringency treatments applied.

Finally, our study demonstrated the usefulness of Hill numbers for comparisons between data sets with large differences in diversity. With the application of Hill numbers, comparability of delivered data can be reached and are highly needed for decision processes in policy making, for example to generalize and prioritize between individual sites or programs. We suggest that the use of Hill numbers and an appropriate stringency treatment is especially relevant for biodiversity monitoring and bioassessment that are mainly interested in taxon richness and less on indices (e.g., Altermatt, Seymour, & Martinez, 2013; Wüthrich & Altermatt, 2019). The intermediate stringency treatment of consistent appearance in at least two filter replicates (i.e., *relaxed*) possibly balances best between the detection of rare ZOTUs and the inflation of diversity measures through sequencing errors, similar to the findings in Alberdi et al. (2018). Regular application of Hill numbers will also advance and unify the field of eDNA metabarcoding to increase comparability across studies and promote the development of best practices.

## Acknowledgements

Data analyzed in this paper were generated in collaboration with the Genetic Diversity Centre (GDC), ETH Zurich. We would like to thank Chelsea J Little, Roman Alther, Emanuel A Fronhofer, Isabelle Gounand, Eric Harvey, Samuel Hürlemann, Simon Flückiger, and Sereina Gut for help in the field and in the lab, and Jeanine Brantschen, Lynn Govaert, and Rosetta Blackman for commenting on the manuscript. Funding is from the Swiss National Science Foundation Grants No PP00P3 179089 and 31003A 173074, the Velux Foundation, and the University of Zurich Research Priority Program “URPP Global Change and Biodiversity” (to FA). All authors declare that there is no conflict of interest regarding the publication of this article.

## Data Accessibility Statement

The sequencing data is available on the European Nucleotide Archive under the following study accession numbers (secondary accession number): PRJEB31920 (ERP114535) and PRJEB33506 (ERP116301).

## Author Contributions

EM and FA designed the study. EM and FA conducted experimental work. EM conducted all laboratory work. EM and JCW performed data analysis. EM wrote the first manuscript draft. EM, JCW, and FA revised the draft.

## References

Adrian-Kalchhauser, I., & Burkhardt-Holm, P. (2016). An eDNA Assay to Monitor a Globally Invasive Fish Species from Flowing Freshwater. PLoS One, 11 (1), e0147558. doi: 10.1371/journal.pone.0147558

Ainsworth, T. D., Krause, L., Bridge, T., Torda, G., Raina, J. B., Zakrzewski, M., … Leggat, W. (2015). The coral core microbiome identifies rare bacterial taxa as ubiquitous endosymbionts. ISME Journal, 9 (10), 2261–2274. doi: 10.1038/ismej.2015.39

Alberdi, A., Aizpurua, O., Gilbert, M. T. P., & Bohmann, K. (2018). Scrutinizing key steps for reliable metabarcoding of environmental samples. Methods in Ecology and Evolution, 9 (1), 134–147. doi: 10.1111/2041-210X.12849

Alberdi, A., & Gilbert, M. T. P. (2019a). A guide to the application of Hill numbers to DNA-based diversity analyses. Molecular Ecology Resources, 19 (4), 804–817. doi: 10.1111/1755-0998.13014

Alberdi, A., & Gilbert, M. T. P. (2019b). hilldiv: an R package for the integral analysis of diversity based on Hill numbers. bioRxiv, 545665.

Altermatt, F., Little, C. J., Mächler, E., Wang, S., Zhang, X., & Blackman, R. C. (2020). Uncovering the complete biodiversity structure in spatial networks–the example of riverine systems. Oikos.

Altermatt, F., Seymour, M., & Martinez, N. (2013). River network properties shape *α*-diversity and community similarity patterns of aquatic insect communities across major drainage basins. Journal of Biogeography, 40 (12), 2249–2260.

Andrews, S. (2015). FASTQC A Quality Control tool for High Throughput Sequence Data. Babraham Institute.

Apothéloz-Perret-Gentil, L., Cordonier, A., Straub, F., Iseli, J., Esling, P., & Pawlowski, J. (2017). Taxonomy-free molecular diatom index for high-throughput eDNA biomonitoring. Molecular ecology resources, 17 (6), 1231–1242.

Bálint, M., Pfenninger, M., Grossart, H. P., Taberlet, P., Vellend, M., Leibold, M. A., … Bowler, D. (2018). Environmental DNA Time Series in Ecology. Trends Ecol Evol. doi: 10.1016/j.tree.2018.09.003

Barbour, M. T., Gerritsen, J., Snyder, B. D., & Stribling, J. B. (1999). Rapid bioassessment protocols for use in streams and wadeable rivers. USEPA, Washington.

BDM Coordination Office. (2009). The state of biodiversity in Switzerland. Overview of the findings of Biodiversity Monitoring Switzerland (BDM) as of May 2009 (Vol. 0911). Biodiversity Monitoring in Switzerland (BDM) Coordination Office, Federal Office for the Environment (BAFU), Bern.

BDM Coordination Office. (2014). Biodiversitätsmonitoring Schweiz BDM. Beschreibung der Methoden und Indikatoren (Vol. 1410). Bundesamt für Umwelt, Bern.

Beentjes, K. K., Speksnijder, A. G. C. L., Schilthuizen, M., Schaub, B. E. M., & van der Hoorn, B. B. (2018). The influence of macroinvertebrate abundance on the assessment of freshwater quality in The Netherlands. Metabarcoding and Metagenomics, 2. doi: 10.3897/mbmg.2.26744

Blaxter, M., Mann, J., Chapman, T., Thomas, F., Whitton, C., Floyd, R., & Abebe, E. (2005). Defining operational taxonomic units using DNA barcode data. Philosophical Transactions of the Royal Society B: Biological Sciences, 360 (1462), 1935–1943.

Bohmann, K., Evans, A., Gilbert, M. T. P., Carvalho, G. R., Creer, S., Knapp, M., … De Bruyn, M. (2014). Environmental DNA for wildlife biology and biodiversity monitoring. Trends in ecology & evolution, 29 (6), 358–367.

Bokulich, N. A., Subramanian, S., Faith, J. J., Gevers, D., Gordon, J. I., Knight, R., … Caporaso, J. G. (2013). Quality-filtering vastly improves diversity estimates from Illumina amplicon sequencing. Nature Methods, 10 (1), 57–59. doi: 10.1038/nmeth.2276

Buchner, D., Beermann, A. J., Laini, A., Rolauffs, P., Vitecek, S., Hering, D., & Leese, F. (2019). Analysis of 13,312 benthic invertebrate samples from german streams reveals minor deviations in ecological status class between abundance and presence/absence data. PloS one, 14 (12).

Chao, A., Chiu, C.-H., & Jost, L. (2014). Unifying species diversity, phylogenetic diversity, functional diversity, and related similarity and differentiation measures through Hill numbers. Annual review of ecology, evolution, and systematics, 45, 297–324.

Chao, A., & Jost, L. (2012). Coverage-based rarefaction and extrapolation: standardizing samples by completeness rather than size. Ecology, 93 (12), 2533–2547.

Chase, J. M., McGill, B. J., McGlinn, D. J., May, F., Blowes, S. A., Xiao, X., … Gotelli, N. J. (2018). Embracing scale-dependence to achieve a deeper understanding of biodiversity and its change across communities. Ecology Letters, 21 (11), 1737–1751.

Colwell, R. K., Chao, A., Gotelli, N. J., Lin, S.-Y., Mao, C. X., Chazdon, R. L., & Longino, J. T. (2012). Models and estimators linking individual-based and sample-based rarefaction, extrapolation and comparison of assemblages. Journal of plant ecology, 5 (1), 3–21.

Colwell, R. K., & Coddington, J. A. (1994). Estimating terrestrial biodiversity through extrapolation. Philosophical Transactions of the Royal Society of London. Series B: Biological Sciences, 345 (1311), 101–118.

Czechowski, P., Stevens, M. I., Madden, C., & Weinstein, P. (2020). Steps towards a more efficient use of chironomids as bioindicators for freshwater bioassessment: Exploiting edna and other genetic tools. Ecological Indicators, 110, 105868.

Deiner, K., Bik, H. M., Mächler, E., Seymour, M., Lacoursiere-Roussel, A., Altermatt, F., … Bernatchez, L. (2017). Environmental DNA metabarcoding: Transforming how we survey animal and plant communities. Mol Ecol, 26 (21), 5872–5895. doi: 10.1111/mec.14350

Deiner, K., Fronhofer, E. A., Mächler, E., Walser, J. C., & Altermatt, F. (2016). Environmental DNA reveals that rivers are conveyer belts of biodiversity information. Nat Commun, 7, 12544. doi: 10.1038/ncomms12544

Deiner, K., Walser, J. C., Mächler, E., & Altermatt, F. (2015). Choice of capture and extraction methods affect detection of freshwater biodiversity from environmental DNA. Biological Conservation, 183, 53–63. doi: 10.1016/j.biocon.2014.11.018

Douglas, W. Y., Ji, Y., Emerson, B. C., Wang, X., Ye, C., Yang, C., & Ding, Z. (2012). Biodiversity soup: metabarcoding of arthropods for rapid biodiversity assessment and biomonitoring. Methods in Ecology and Evolution, 3 (4), 613–623.

Dudgeon, D., Arthington, A. H., Gessner, M. O., Kawabata, Z.-I., Knowler, D. J., Lévêque, C., … Sullivan, C. A. (2006). Freshwater biodiversity: importance, threats, status and conservation challenges. Biological Reviews, 81 (2), 163–182. doi: 10.1017/S1464793105006950

Edgar, R. C. (2016). UNOISE2: improved error-correction for Illumina 16S and ITS amplicon sequencing. bioRxiv, 81257. doi: 10.1101/081257

Elbrecht, V., & Leese, F. (2015). Can DNA-based ecosystem assessments quantify species abundance? Testing primer bias and biomass—sequence relationships with an innovative metabarcoding protocol. PloS one, 10 (7), e0130324.

Elbrecht, V., Vamos, E. E., Meissner, K., Aroviita, J., & Leese, F. (2017). Assessing strengths and weaknesses of DNA metabarcoding-based macroinvertebrate identification for routine stream monitoring. Methods in Ecology and Evolution, 8 (10), 1265–1275. doi: 10.1111/2041-210X.12789

Eren, A. M., Sogin, M. L., Morrison, H. G., Vineis, J. H., Fisher, J. C., Newton, R. J., & McLellan, S. L. (2015). A single genus in the gut microbiome reflects host preference and specificity. The ISME Journal, 9 (1), 90–100. doi: 10.1038/ismej.2014.97

Evans, N. T., Li, Y., Renshaw, M. A., Olds, B. P., Deiner, K., Turner, C. R., … Pfrender, M. E. (2017). Fish community assessment with eDNA metabarcoding: effects of sampling design and bioinformatic filtering. Canadian Journal of Fisheries and Aquatic Sciences, 74 (9), 1362–1374.

Ferraro, S. P., Cole, F. A., DeBen, W. A., & Swartz, R. C. (1989). Power-cost efficiency of eight macrobenthic sampling schemes in Puget Sound, Washington, USA. Canadian Journal of Fisheries and Aquatic Sciences, 46 (12), 2157–2165.

Geller, J., Meyer, C., Parker, M., & Hawk, H. (2013). Redesign of PCR primers for mitochondrial cytochrome c oxidase subunit I for marine invertebrates and application in all-taxa biotic surveys. Mol Ecol Resour, 13 (5), 851–861. doi: 10.1111/1755-0998.12138

Giraudoux, P. (2018). pgirmess: Spatial Analysis and Data Mining for Field Ecologists. R package version, 361 (1.6), 9.

Good, I. J. (1953). The population frequencies of species and the estimation of population parameters. Biometrika, 40 (3-4), 237–264.

Good, I. J. (2000). Turing’s anticipation of empirical bayes in connection with the cryptanalysis of the naval enigma. Journal of Statistical Computation and Simulation, 66 (2), 101–111.

Gotelli, N. J. (2001). A primer of ecology. Sunderland, Massachusetts. Sinauer Associates, Inc.

Gotelli, N. J., & Colwell, R. K. (2011). Estimating species richness. Biological diversity: frontiers in measurement and assessment, 12, 39–54.

Grooten, M., Almond, R., et al. (2018). Aiming higher. Living planet report 2018.

Haase, P., Lohse, S., Pauls, S., Schindehütte, K., Sundermann, A., Rolauffs, P., & Hering, D. (2004). Assessing streams in germany with benthic invertebrates: development of a practical standardised protocol for macroinvertebrate sampling and sorting. Limnologica, 34 (4), 349–365.

Hänfling, B., Lawson Handley, L., Read, D. S., Hahn, C., Li, J., Nichols, P., … Winfield, I. J. (2016). Environmental DNA metabarcoding of lake fish communities reflects long-term data from established survey methods. Molecular Ecology.

Hering, D., Johnson, R. K., Kramm, S., Schmutz, S., Szoszkiewicz, K., & Verdonschot, P. F. (2006). Assessment of european streams with diatoms, macrophytes, macroinvertebrates and fish: a comparative metric-based analysis of organism response to stress. Freshwater Biology, 51 (9), 1757–1785.

Hill, M. O. (1973). Diversity and evenness: a unifying notation and its consequences. Ecology, 54 (2), 427–432.

Hsieh, T., Ma, K., & Chao, A. (2016). iNEXT: an R package for rarefaction and extrapolation of species diversity (Hill numbers). Methods in Ecology and Evolution, 7 (12), 1451–1456.

Johnson, R. K., Furse, M. T., Hering, D., & Sandin, L. (2007). Ecological relationships between stream communities and spatial scale: implications for designing catchment-level monitoring programmes. Freshwater Biology, 52 (5), 939–958.

Jost, L. (2007). Partitioning diversity into independent alpha and beta components. Ecology, 88 (10), 2427–2439.

Kang, S., Rodrigues, J. L., Ng, J. P., & Gentry, T. J. (2016). Hill number as a bacterial diversity measure framework with high-throughput sequence data. Scientific reports, 6, 38263.

Kelly, M., Juggins, S., Guthrie, R., Pritchard, S., Jamieson, J., Rippey, B., … Yallop, M. (2008). Assessment of ecological status in UK rivers using diatoms. Freshwater Biology, 53 (2), 403–422.

Leese, F., Bouchez, A., Abarenkov, K., Altermatt, F., Borja, Á., Bruce, K., … others (2018). Why we need sustainable networks bridging countries, disciplines, cultures and generations for aquatic biomonitoring 2.0: a perspective derived from the DNAqua-Net COST action. In Advances in ecological research (Vol. 58, pp. 63–99). Elsevier.

Leray, M., & Knowlton, N. (2017). Random sampling causes the low reproducibility of rare eukaryotic OTUs in Illumina COI metabarcoding. PeerJ, 5, e3006. doi: 10.7717/peerj.3006

Leray, M., Yang, J. Y., Meyer, C. P., Mills, S. C., Agudelo, N., Ranwez, V., … Machida, R. J. (2013). A new versatile primer set targeting a short fragment of the mitochondrial COI region for metabarcoding metazoan diversity: application for characterizing coral reef fish gut contents. Frontiers in zoology, 10 (1), 34.

Macher, J.-N., & Leese, F. (2017). Environmental DNA metabarcoding of rivers: Not all eDNA is everywhere, and not all the time. bioRxiv. doi: 10.1101/164046

Macher, J.-N., Vivancos, A., Piggott, J. J., Centeno, F. C., Matthaei, C. D., & Leese, F. (2018). Comparison of environmental DNA and bulk-sample metabarcoding using highly degenerate cytochrome c oxidase I primers. Molecular Ecology Resources, 18 (6), 1456–1468. doi: 10.1111/1755-0998.12940

Mächler, E., Deiner, K., Steinmann, P., & Altermatt, F. (2014). Utility of Environmental DNA for Monitoring Rare and Indicator Macroinvertebrate Species. Freshwater Science, 33 (4), 1174–1183. doi: 10.1086/678128

Mächler, E., Little, C. J., Wüthrich, R., Alther, R., Fronhofer, E. A., Gounand, I., … Altermatt, F. (2019, sep). Assessing different components of diversity across a river network using eDNA. Environmental DNA, 1 (3), 290–301. doi: 10.1002/edn3.33

Manley, P. N., & Van Horne, B. (2006). The multiple species inventory and monitoring protocol: a population, community, and biodiversity monitoring solution for national forest system lands (Vol. 42).

McGee, K. M., Robinson, C., & Hajibabaei, M. (2019). Gaps in DNA-based Biomonitoring Across the Globe. Frontiers in Ecology and Evolution, 7, 337.

McKee, A. M., Spear, S. F., & Pierson, T. W. (2015). The effect of dilution and the use of a post-extraction nucleic acid purification column on the accuracy, precision, and inhibition of environmental DNA samples. Biological Conservation, 183, 70–76. doi: 10.1016/j.biocon.2014.11.031

McMurdie, P. J., & Holmes, S. (2013). phyloseq: an R package for reproducible interactive analysis and graphics of microbiome census data. PloS one, 8 (4), e61217.

McMurdie, P. J., & Holmes, S. (2014). Waste Not, Want Not: Why Rarefying Microbiome Data Is Inadmissible. PLOS Computational Biology, 10 (4), e1003531.

Murray, D. C., Coghlan, M. L., & Bunce, M. (2015). From benchtop to desktop: important considerations when designing amplicon sequencing workflows. PLoS One, 10 (4).

Pawlowski, J., Kelly-Quinn, M., Altermatt, F., Apothéloz-Perret-Gentil, L., Beja, P., Boggero, A., … others (2018). The future of biotic indices in the ecogenomic era: Integrating (e)DNA metabarcoding in biological assessment of aquatic ecosystems. Science of the Total Environment, 637, 1295–1310.

Porter, T. M., & Hajibabaei, M. (2018). Scaling up: A guide to high-throughput genomic approaches for biodiversity analysis. Molecular ecology, 27 (2), 313–338.

R Core Team. (2018). R: A language and environment for statistical computing.

Simberloff, D. (1978). Use of rarefaction and related methods in ecology. In Biological data in water pollution assessment: quantitative and statistical analyses. ASTM International.

Taberlet, P., Bonin, A., Zinger, L., & Coissac, E. (2018). Environmental DNA: For Biodiversity Research and Monitoring. Oxford University Press.

Taberlet, P., Coissac, E., Hajibabaei, M., & Riesenberg, L. H. (2012). Environmental DNA. Molecular Ecology, 21 (8), 1789–1793.

Tachet, H., Richoux, P., Bournaud, M., & Usseglio-Polatera, P. (2010). Invertébrés d’eau douce: systématique, biologie, écologie (Vol. 15). CNRS éditions Paris.

Visco, J. A., Apothéloz-Perret-Gentil, L., Cordonier, A., Esling, P., Pillet, L., & Pawlowski, J. (2015). Environmental monitoring: inferring the diatom index from Next-Generation Sequencing data. Environmental science & technology, 49 (13), 7597–7605.

Vivien, R., Apothéloz-Perret-Gentil, L., Pawlowski, J., Werner, I., & Ferrari, B. J. (2019). Testing different (e)DNA metabarcoding approaches to assess aquatic oligochaete diversity and the biological quality of sediments. Ecological Indicators, 106, 105453.

Vörösmarty, C. J., McIntyre, P. B., Gessner, M. O., Dudgeon, D., Prusevich, A., Green, P., … Liermann, C. R. (2010). Global threats to human water security and river biodiversity. Nature, 467 (7315), 555–561.

Weigand, H., Beermann, A. J., Čiampor, F., Costa, F. O., Csabai, Z., Duarte, S., … Ekrem, T. (2019). DNA barcode reference libraries for the monitoring of aquatic biota in Europe: Gap-analysis and recommendations for future work. Science of The Total Environment, 678, 499–524. doi: https://doi.org/10.1016/j.scitotenv.2019.04.247

Wüthrich, R., & Altermatt, F. (2019). Aquatische Monitoringprogramme NAWA und BDM. Synergien, Strategien und Visionen. Bundesamt für Umwelt (BAFU).

Yamamoto, S., Masuda, R., Sato, Y., Sado, T., Araki, H., Kondoh, M., … Miya, M. (2017). Environmental DNA metabarcoding reveals local fish communities in a species-rich coastal sea. Sci Rep, 7, 40368. doi: 10.1038/srep40368

Zhan, A., Hulák, M., Sylvester, F., Huang, X., Adebayo, A. A., Abbott, C. L., … MacIsaac, H. J. (2013). High sensitivity of 454 pyrosequencing for detection of rare species in aquatic communities. Methods in Ecology and Evolution, 4 (6), 558–565.

