## Supplementary file for "Decision making and best practices for taxonomy-free eDNA metabarcoding in biomonitoring using Hill numbers"

### Contents

**Table S1 - Diversity partitioning** **2**

**Table S2 - Results Kruskal-Wallis test for beta diversity** **3**

**Figure S1 - ZOTU abundances between runs** **4**

**Figure S2 - Rarefaction curves** **5**

**Figure S3 - Beta diversity** **6**

### Table S1 - Diversity partitioning

**Table S1:** Diversity partitioning of the five data sets for Hill numbers of order 0 to 2, in steps of 0.5. Values are rounded to two digits.

| Order | Stringency treatment | Alpha | Gamma | Beta |
|---|---|---|---|---|
| 0 | Additive | 1494.29 | 11129.00 | 7.45 |
| | Relaxed | 585.82 | 5948.00 | 10.15 |
| | Strict | 265.05 | 3167.00 | 11.95 |
| | Threshold | 569.14 | 1361.00 | 2.39 |
| | Rarefied | 1301.04 | 11102.00 | 8.53 |
| 0.5 | Additive | 323.48 | 2216.44 | 6.85 |
| | Relaxed | 173.07 | 1196.78 | 6.92 |
| | Strict | 95.71 | 738.58 | 7.72 |
| | Threshold | 161.98 | 692.34 | 4.27 |
| | Rarefied | 313.29 | 2209.38 | 7.05 |
| 1 | Additive | 76.81 | 503.49 | 6.56 |
| | Relaxed | 53.51 | 342.09 | 6.39 |
| | Strict | 36.31 | 249.69 | 6.88 |
| | Threshold | 53.73 | 322.86 | 6.01 |
| | Rarefied | 76.58 | 503.09 | 6.57 |
| 1.5 | Additive | 27.47 | 213.10 | 7.76 |
| | Relaxed | 22.09 | 164.98 | 7.47 |
| | Strict | 17.27 | 131.50 | 7.61 |
| | Threshold | 22.19 | 173.61 | 7.58 |
| | Rarefied | 27.46 | 213.03 | 7.76 |
| 2 | Additive | 14.44 | 131.23 | 9.09 |
| | Relaxed | 12.37 | 107.21 | 8.67 |
| | Strict | 10.44 | 88.94 | 8.52 |
| | Threshold | 12.88 | 113.98 | 8.85 |
| | Rarefied | 14.44 | 131.18 | 9.09 |

### Table S2 - Results Kruskal-Wallis test for beta diversity

**Table S2:** Results of Kruskal-Wallis test on ranks of the beta diversity and the following multiple mean comparisons post-hoc tests based on rank sums. Chi-Squared, degrees of freedom (DF), number of data points (n) and p-values are given. Stringency treatments with the same letter are not significantly different according to multiple comparison post-hoc test after Kruskal-Wallis (p-value = 0.05).

| Kruskal-Wallis test on ranks | | | | | Multiple mean comparisons post-hoc tests | | | | |
|---|---|---|---|---|---|---|---|---|---|
| **Hill order** | **Chi-Squared** | **DF** | **n** | **p-value** | **Additive** | **Relaxed** | **Strict** | **Threshold** | **Rarefied** |
| 0 | 3829.9 | 4 | 7700 | $< 0.001$ | a | b | c | b | e |
| 1 | 115.4 | 4 | 7700 | $< 0.001$ | a | a | b | c | a |
| 2 | 3.8 | 4 | 7700 | 0.43 | NA | NA | NA | NA | NA |

### Figure S1 - ZOTU abundances between runs

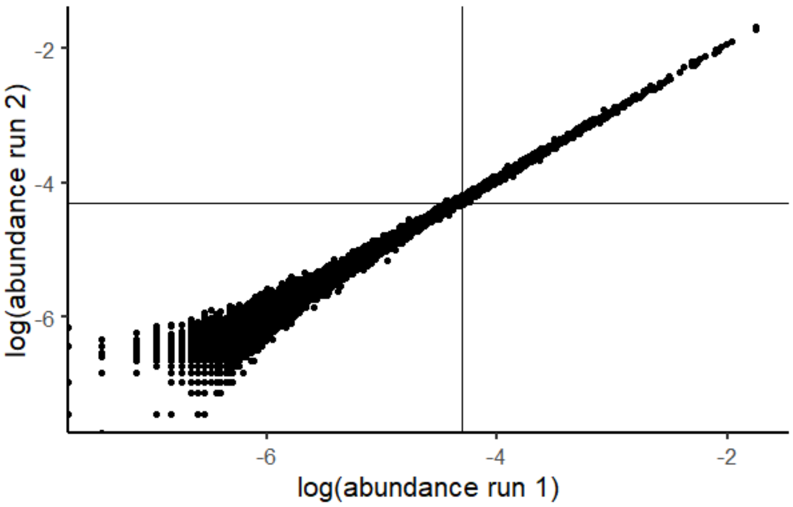


**Fig. S1:** To estimate consistency between the two runs, we plotted the abundances of the first run against the second run for each ZOTU. The solid lines indicate the chosen cut-off of 0.005%. ZOTUs with a higher abundance than this cut-off have highly similar abundances in the two runs

### Figure S2 - Rarefaction curves

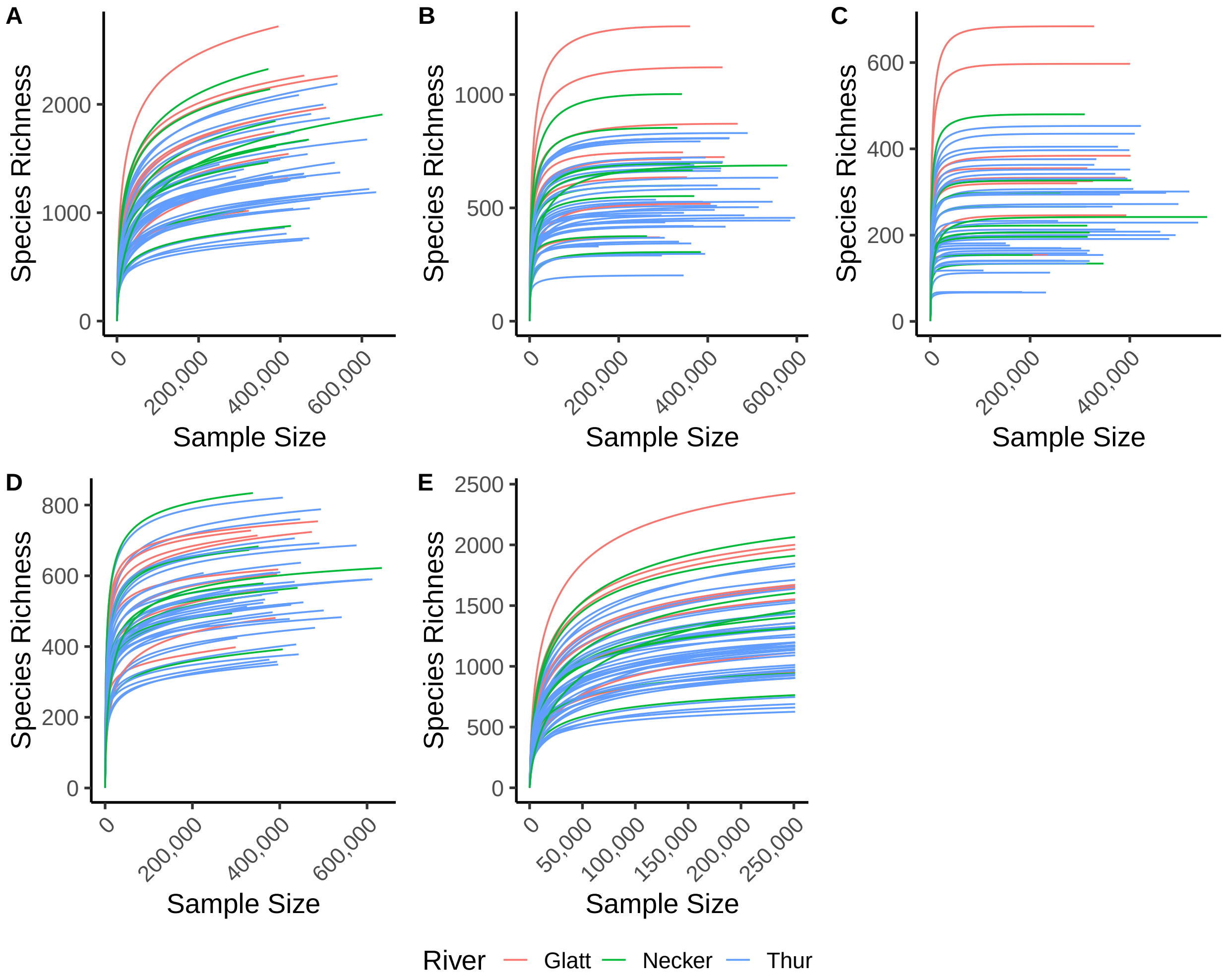


**Fig. S2:** Rarefaction curves for the individual sites for the specific stringency treatments. (A) *additive*, (B) *relaxed*, (C) *strict*, (D) *threshold*, and (E) *rarefied*.

### Figure S3 - Beta diversity

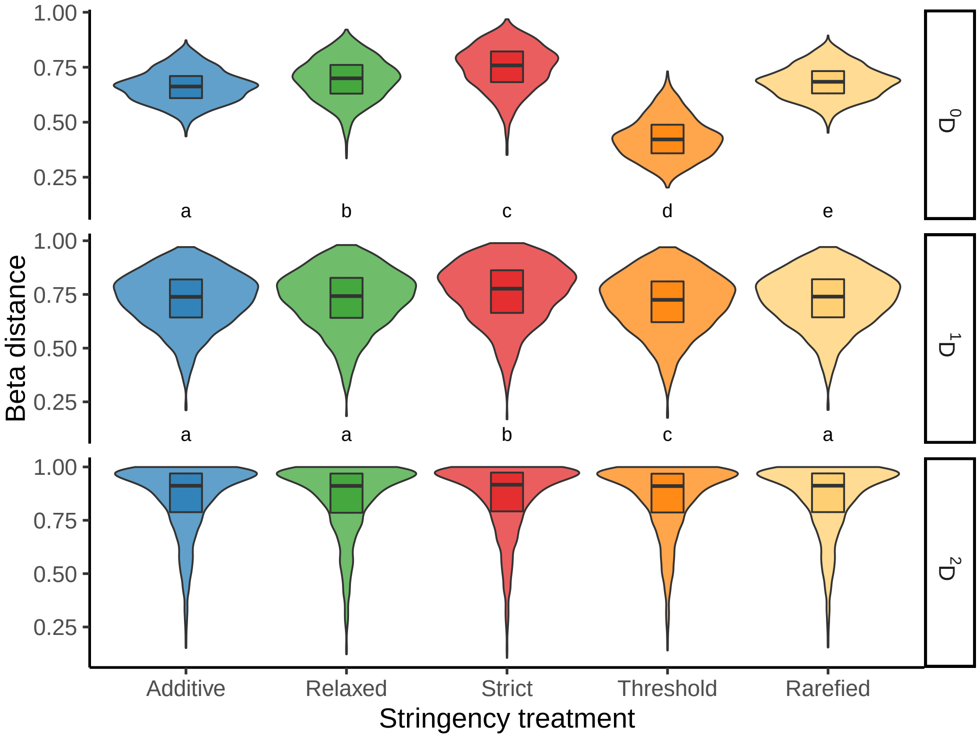


**Fig. S3:** Beta distance between the individual sites of the five different stringency treatments for the Hill order $q = 0$ (i.e., the Jaccard index, upper panel), $q = 1$ (middle panel), and $q = 2$ (i.e., Morisita-Horn index, lower panel). Stringency treatments with dissimilar letters are significantly different according to multiple mean comparison test after Kruskal-Wallis applied separately to the individual Hill number orders.
